# An evidence assessment tool for ecosystem services and conservation studies

**DOI:** 10.1101/010140

**Authors:** Anne-Christine Mupepele, Jessica C. Walsh, William J. Sutherland, Carsten F. Dormann

## Abstract

Reliability of scientific findings is important, especially if they directly impact decision making, such as in environmental management. In the 1990s, assessments of reliability in the medical field resulted in the development of evidence-based practice. Ten years later, evidence-based practice was translated into conservation, but so far no guidelines exist on how to assess the evidence of individual studies. Assessing the evidence of individual studies is essential to appropriately identify and summarize the confidence in research findings. We develop a tool to assess the strength of evidence of ecosystem services and conservation studies. This tool consists of (1) a hierarchy of evidence, based on the experimental design of studies and (2) a critical-appraisal checklist that identifies the quality of research implementation. The application is illustrated with 13 examples and we suggest further steps required to move towards more evidence-based environmental management.

## In a nutshell

- Human’s life depends on nature, biodiversity and their related ecosystem services and it is essential to manage natural ecosystems in a sustainable way.
- Decisions taken in the context of environmental management need to be based on sound knowledge and reliable information.
- We introduce an evidence assessment tool to identify the reliability of environmental research.
- This evidence assessment tool is based on a hierarchical ranking of study designs and a study quality checklist.
- Identifying the reliability of environmental research is the first step towards more effective decision making.

Conservation and ecosystem services studies are important scientific sources for decision-makers seeking advice on environmental management (Daily and Matson, 2008; Kareiva and Marvier, 2012). These study results potentially influence actions and it is therefore crucial to assess transparently the reliability of current research and its recommendations (Pullin and Knight, 2003; Boyd, 2013).

Evidence-based practice was introduced in the medical field (Sackett *et al.*, 1996; GRADE Working Group, 2004; OCEBM Levels of Evidence Working Group, 2011, Cochrane Collaboration - www.cochrane.org) aiming to assess the reliability of scientific statements and identify the best available information to answer a question of interest (Sackett *et al.*, 1996). In conservation, evidence-based practice was first mentioned 15 years ago (Sutherland, 2000; Pullin and Knight, 2001). Today, the Collaboration for Environmental Evidence (www.environmentalevidence.org) fosters the creation of systematic reviews to collate the strongest possible evidence (Petrokofsky *et al.*, 2011; Bowler *et al.*, 2012; Collaboration for Environmental Evidence, 2013, see also *Journal for Environmental Evidence*), together with the Conservation Evidence (www.conservationevidence.org), which focuses on the development of summaries and guidelines, and the communication of evidence to practitioners (Sutherland *et al.*, 2012; Dicks *et al.*, 2014).

Alongside these ongoing attempts we identify the need for a clear hierarchical ranking of study designs used in ecosystem services and conservation in order to evaluate available information. Here we first discuss the terminology of evidence-based practice, to ensure that scientists and practitioner can communicate effectively across the disciplines and backgrounds. Next, we introduce a new evidence hierarchy that ranks scientific study designs in ecosystem services and conservation, extending a previous proposal by Pullin and Knight (2003). A quality checklist will support the appraisal of the study design and increase reproducibility. Finally, we illustrate the application of the tool with 13 case studies, and specify the relevance of evidence-based practice for different user groups.

## Current use of evidence-based practice in environmental management

A common application in evidence-based practice is a **systematic review**. It summarises the knowledge available for a specific question using systematic and explicit methods to identify and select relevant research (Collaboration for Environmental Evidence, 2013). Another approach to evidence-based practice are **summaries**. Summaries do not focus on a specific question but bring together information from a much broader topic, e.g. from a whole animal group (Dicks *et al.*, 2014), such as bees (Dicks *et al.*, 2010). They assess the evidence of various possible interventions, but do not currently appraise the quality of each study included or give specific recommendations. Summaries can serve as a foundation to develop **guidelines**. These ‘best practice guides’ give recommendations on conservation strategies and ecosystem services assessment tools. They are based on the collection of scientific evidence summarized and judged by a group of experts (Graham *et al.*, 2011; Sutherland *et al.*, 2015).

Systematic reviews and summaries compile individual studies and therefore require the evaluation of the evidence at the level of the individual study. In systematic reviews this is typically mentioned as one step of the critical appraisal. However, to date such critical appraisal is often implicit, based on criteria varying for every systematic review (Collaboration for Environmental Evidence, 2013). The instruments introduced here provide a clear appraisal guideline with an evidence assessment tool to score the reliability of individual studies.

## Evidence assessment tool

The terminology used around evidence-based practice is diverse and not always consistently used. However, a well-defined terminology is essential for effective communication between practitioners and scientists. According to the Oxford Dictionaries, **evidence** is ‘the available body of […] information indicating whether a belief or proposition is true or valid’ (www.oxforddictionaries.com/definition/english/evidence). Evidence describes the knowledge behind a statement and expresses how solid our recommendations are (see also Higgs and Jones 2000, p.311; Rychetnik *et al.* 2001; Lohr 2004; Pullin and Knight 2005). The **strength of evidence** reflects the reliability of information and we can identify whether a statement is based on **strong or weak evidence**, i.e. very reliable or hardly reliable. The application of evidence-based practice requires the identification and use of the best available evidence underlying a statement. To ensure reproducibility, the collation and appraisal of the best-available evidence should follow explicit criteria.

### 1. Setting question and context

The formulation of a clear research question and the purpose of investigation is highly emphasized throughout the evidence literature (Higgins and Green, 2011; Collaboration for Environmental Evidence, 2013, p.20-23). Questions should specify *which* ecosystem service, species or aspect of biodiversity will be investigated in *which* system, as this will help to determine the external validity of the answer provided in a study.

We further recommend to determine the focus of the question, as either ‘quantification’, ‘valuation’, ‘management’ or ‘governance’. **Quantification** studies measure the amount of an ecosystem service, species abundance, biodiversity or other conservation targets. Examples include estimating species abundance, the quantity of carbon sequestrated or the number of flowers pollinated. Measures can be taken in absolute units or relative to another system. **Valuation** studies assess the societal value of ecosystem services. The most common way is monetary valuation. Other possibilities are in relation to a reference system or on a ranked scale (high, middle, low value). **Management** is the treatment designed to improve or benefit specific ecosystem services, target species, biodiversity or other conservation aspects. For example: leaving dead wood in forests to increase biodiversity or reducing agricultural fertiliser to decrease nearby lake eutrophication. **Governance** is seen as the strategy or policy to steer a management intervention, such as REDD (Reducing Emissions from Deforestation and forest Degradation), which aims to encourage forest protection and reforestation (Kenward *et al.*, 2011). The tools used by policy makers include incentives (subsidiaries) or penalties (law/tax) (see also Bevir, 2012). When the effectiveness of management and governance strategies is determined, evidence-based quantification or valuation is required to measure the outcome of the management or governance intervention. Acuña *et al.* (2013), for example, uses valuation methods to determine success or failure of a management strategy while Walsh *et al.* (2012) quantifies malleefowl abundance through monitoring survey data to assess the management impact of fox baiting on malleefowl. The distinction of four different foci is essential to assess the whole range of environmental management.

We have described how to set the context of questions that can be useful in environmental management. Once the question has been determined, and the investigation carried out, the strength of the resulting evidence should be assessed (Fig. 1).

**Figure 1.**
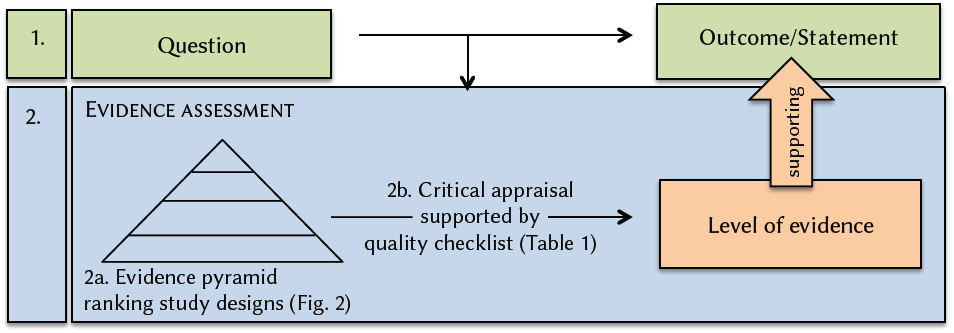
Schematic procedure of evidence-based practice: 1. Identification of study question and results. 2. Assessing the strength of evidence supporting the result, with help of a evidence hierarchy and a quality checklist.

### 2. Evidence assessment

The reliability of a study is characterized by its study design and the quality of its implementation. Both are evaluated in the evidence assessment.

#### 2a. Evidence hierarchy

The study design refers to the set-up of the investigation, e.g. controlled or observational design (GRADE Working Group, 2004). These study designs are not equally compelling with respect to inferring causality. Differences in study designs typically translate into weak or strong evidence. To identify the reliability of a study, study designs can be ranked hierarchically according to a level-of-evidence scale, hence forth the evidence hierarchy (Fig. 2).

**Figure 2.**
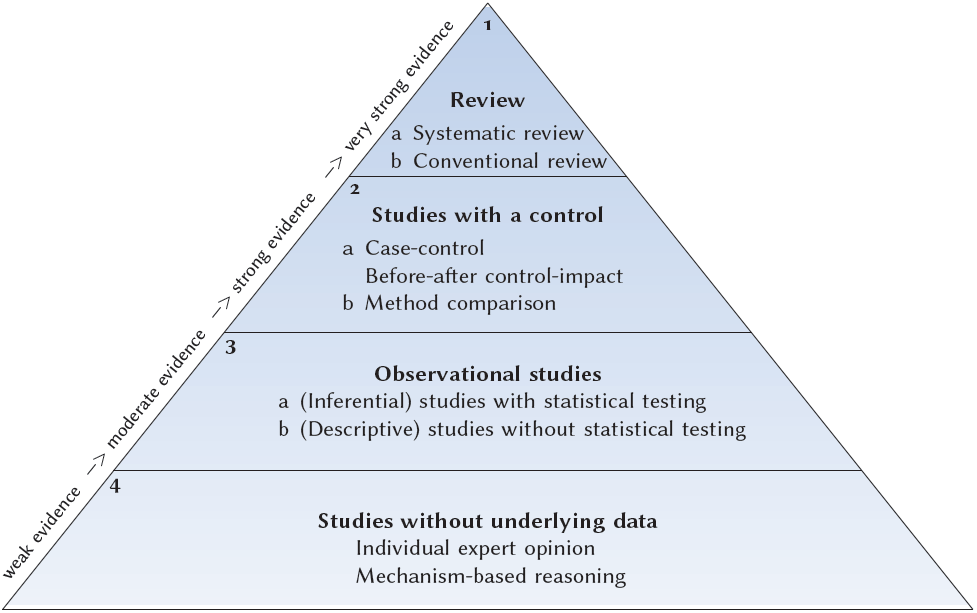
Level-of-evidence (LoE) hierarchy ranking study designs according to their evidence. LoE1 - LoE4 with internally ranked sublevels a and b.

**Systematic reviews (LoE1a)** are at the top of the evidence hierarchy and provide the most reliable information. They summarize all information collated in several individual studies, have an *a priori* protocol on design and procedure, and are conducted according to strict guidelines (e.g. Collaboration for Environmental Evidence, 2013). If possible, they ideally include quantitative measures, at best a meta-analysis. Other more **conventional reviews (LoE1b)** may also include quantitative analysis or are purely qualitative. Both types of review summarize the findings of several studies, but systematic reviews assess the completeness and reproducibility more carefully and strive to reduce bias by having transparent, thorough, pre-approved methods (Higgins and Green, 2011).

The necessary condition for any review is that appropriate individual studies are available. The most reliable individual study design is **a study with a reference/control (LoE2)**. Typically, these are case-control or before-after control-impact studies **(LoE2a)** (Smith *et al.***, 2014**). Another useful approach can be the comparison of different treatments or interventions, for example for the valuation of ecosystem services, where no control exists. Comparing results of different valuation approaches can increase the evidence, if results of both approaches are consistent **(LoE2b)**.

**Observational studies (LoE3)** are individual studies without a control. These include studies employing inferential and correlative statistics, e.g. testing for the influence of environmental variables on the quantity of an ecosystem service **(LoE3a)**. Descriptive studies imply data collection and representation without statistical testing (e.g. data summaries, ordinations, histograms, models with data input). In ecosystem services science and conservation these are often surveys or expert elicitations **(LoE3b)**.

The lowest level of evidence are statements **without underlying data (LoE4)**. These are usually individual expert opinions, often not distinguishable from randomness (Tetlock, 2005). Other statements without underlying data are reasoning based on mechanism and ‘first principles’: *A* works according to a certain mechanism, so we expect *B* to work in the same way. These first principles are not reliable in ecology (Lawton, 1999).

It is important to note that ‘method’ and ‘design’ should not be confused. Methods are the means used to collect or analyse data, e.g. remote sensing, questionnaires, ordination techniques, model types. Design reflects how the study was planned and conducted, e.g. a case-control or observational design (GRADE Working Group, 2004). The same methods can be employed for different underlying designs. Remote sensing for example can be done purely descriptively or with a reference such as ground-truthing or in a ‘before-and-after’ design.

#### 2b. Critical appraisal

Study design alone is an inadequate marker of the strength of evidence (Rychetnik *et al.*, 2001). A study with a strong-evidence design may be poorly conducted. The critical appraisal assesses the implementation of the study design, specifically the methodological quality, the actual realization of the study design and its reporting (Higgins and Green, 2011, section 15.5.2). It identifies the study quality and may lead to a downgrading in the evidence hierarchy. **Quality**, in this context, is the extent to which all aspects of conducting a study can be shown to protect against bias, and inferential error (Lohr, 2004). Qyality checklists should detect bias and inferential error. Combining 30 published quality checklists, we provide the first quality checklist for conservation and ecosystem services (Table 1, WebTable 1), that can be used to comprehensively assess the internal validity of a study, covering questions on data collection, analysis and the presentation of results. The checklist consists of 43 questions, of which some apply only to a specific context, e.g. for reviews or only studies focusing on valuation. All questions answered with ‘yes’ receive one point (or two points for an important question). In the case of non-reported issues, we advise the answer ‘no’ to indicate a deficient reporting quality. The percentage of points received can then help to decide whether to downgrade the level of evidence (Table 2).

**Table 1.**
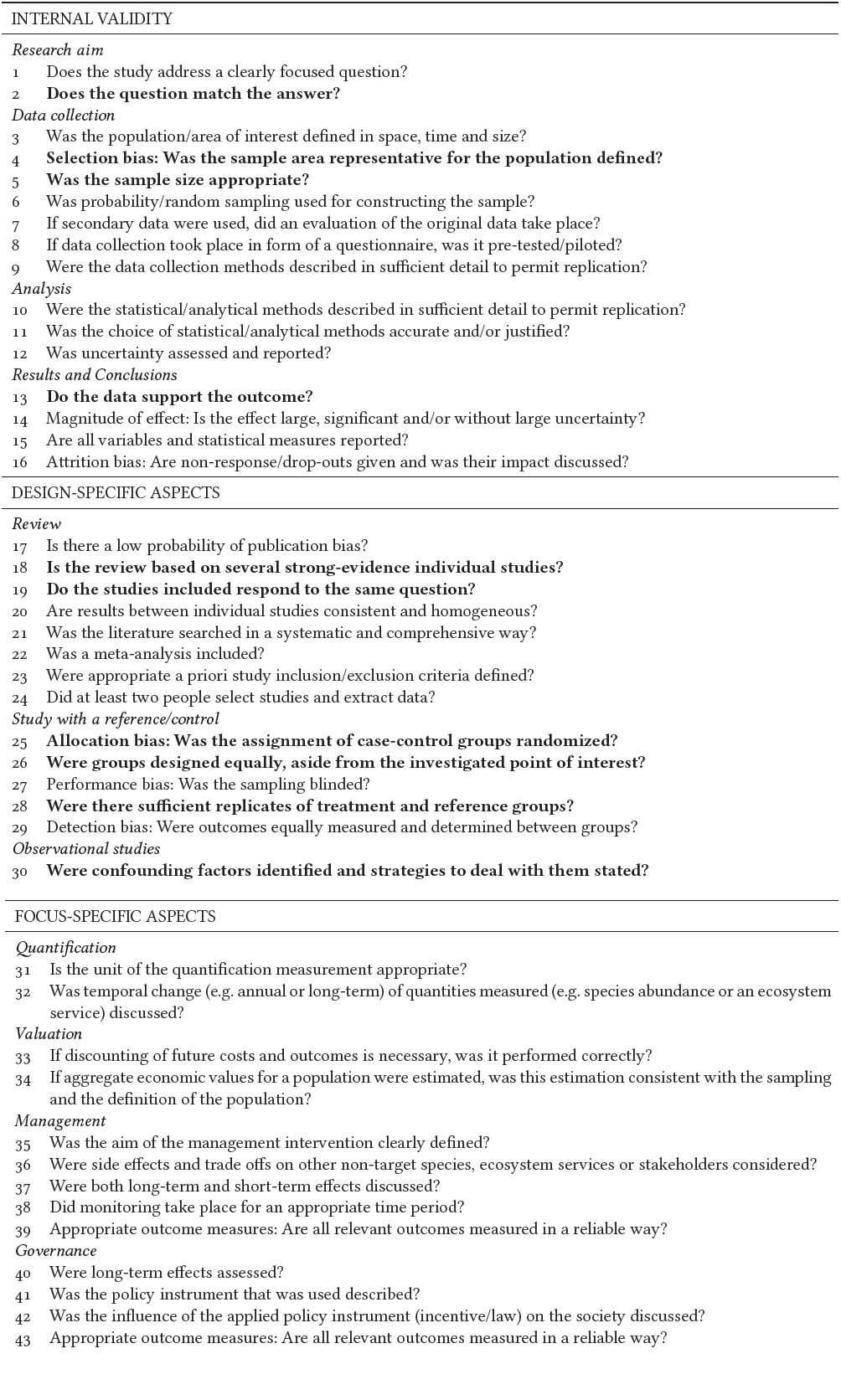
**Qyality checklist** questions. Each question answered with ‘yes’ will receive one point, important aspects (bold type) two points. If a question is not appropriate, it may be left out.

**Table 2.** Downgrading the level of evidence (LoE) according to the percentage of quality points reached

| Percentage of quality points | Downgrading of the LoE |
| --- | --- |
| > 87% of total points | -> no downgrading |
| 75 - 87% of total points | -> downgrading by half a level (e.g. LoE1a to LoE1b) |
| 62 - 74% of total points | -> downgrading by one level (e.g. LoE1a to LoE2a) |
| 50 - 61% of total points | -> downgrading by one and a half levels (e.g. LoE1a to LoE2b) |
| 37 - 49% of total points | -> downgrading by two levels |
| 25 - 36% of total points | -> downgrading by two and a half levels |
| < 24% of total points | -> downgrading by three levels |

Reviews provide information at the highest level of evidence and their critical appraisal is different from other designs, because they are based on studies with weaker evidence (see Table 1: Review). If only studies based on weak evidence were included, then the review should be downgraded, regardless of other quality criteria. A review can be assessed for its quality using our checklist. Furthermore every single study included in the review can also be assessed for its level of evidence, again using the checklist for quality criteria and the evidence hierarchy.

The checklist should make the assessment more transparent, but we are aware that many questions in the checklist can be subjective and depend on the judgement of the assessor (Cohen’s kappa test for two raters: 0.49). We encourage the use of the checklist for an orientation, but we want to emphasise that this procedure can not be fully standardised.

The combination of study design and quality criteria identifies the level of evidence supporting the study result (Fig. 1).

## Application of the evidence assessment tool

The suggested method was applied to assess the evidence of 13 studies (WebTable 2). They were selected to serve as examples and illustrate the applicability of the tool to the whole range of study designs and foci. The first example was a management-related systematic review of Mant *et al.* (2013), conducted according to the guidelines of the Collaboration for Environmental Evidence (2013). They investigated the effect of ‘liming’ rivers or lakes on fish and invertebrate populations. They found that liming increased fish abundances and acid-sensitive invertebrates, but may have a negative impact on the abundance of all invertebrate taxa combined. According to the critical appraisal the study achieved 25 out of 28 points (89%) and it therefore remained at the originally assigned LoE1a, the highest level of evidence.

A second example tackles the question: ‘How does adding dead wood to rivers influence the provision of ecosystem services?’ (Acuña *et al.*, 2013). The authors investigated two ecosystem services (fishing and retention of organic and inorganic matter) in a river-forest ecosystem in Spain and Portugal and studied the effect of this management intervention. Their study design followed a before-after control-impact approach, equivalent to LoE2a. The critical appraisal revealed shortcomings, e.g. no blinding, no randomization and no probability sampling: only 22 out of 31 points (71%) were achieved. The level of evidence was downgraded by one level to LoE3a. We therefore conclude that the statement made by Acuña *et al.* (2013): ‘restoration of natural wood loading in streams increases the ecosystem service provision’ is based on moderate evidence (LoE3a).

We provide further examples in the supporting information, together with the detailed quality checklist filled in for each study (WebTable 2, WebTable 3 and GitHub: https://github.com/biometry/EvidenceAssessmentTool/blob/master/Examples.xlsx). All but one studies revealed quality shortcomings and had to be downgraded. Most were scored as LoE3 or LoE4.

## Relevance for different user groups

In the previous section it was elaborated *how* to assess the strength of evidence for individual studies and reviews. Now we provide a few notes on *who* should use it:

1. **Scientists conducting their own studies** have to be aware of how to achieve strong evidence, particularly during the planning phase. Choosing a study design that provides strong evidence and respects the quality criteria will substantially increase the potential contribution to our knowledge.
2. **Scientists advising decision-makers** should be explicit about the strength of evidence of information they include in their recommendations. Weighting all scientific information equally, or subjectively, runs the risk of overconfidence and bias.
3. **Decision-makers** receiving information from scientists should demand a level-of-evidence statement for the information provided. Alternatively, they can judge themselves the reliability having in mind the assessment for the evidence, although the latter one might be difficult as some scientific training is necessary to identify the study design and evaluate most of the quality questions.
4. **Research funders** should demand scientists to state how they intend to achieve strong evidence results and provide a level-of-evidence statement.
5. We further would like to encourage **consortia, international panels and learned societies**, such as the Intergovernmental Platform on Biodiversity & Ecosystem Services (IPBES), the Ecological Societies (ESA, BES, GFÖ and others), the Society for Conservation Biology (SCB) and the Ecosystem Services Partnership (ESP) to support the development of guidelines (Graham *et al.*, 2011; Sutherland *et al.*, 2015). These guidelines contain recommendations on how to best quantify, value, manage or govern a desired ecosystem service or conservation target. This would give decision-makers more transparent summarised advice, and decrease work load and therefore costs of advisory panels.

## Conclusion

We outlined an evidence assessment tool for ecosystem services and conservation studies, encompassing a hierarchy to judge the available evidence based on study design and a quality checklist to facilitate critical appraisal. We further illustrated with examples how to apply the tool (see also supporting information). Evidence-based practice does not contradict other existing management concepts. It complements these approaches, emphasising that whatever information is used to inform decision should be accompanied by awareness of how reliable this knowledge is.

We are aware of criticism of evidence hierarchies claiming that controlled trials are not always more reliable than observational studies (Petticrew and Roberts, 2003). With our quality checklist we emphasize the critical appraisal to check for an appropriate implementation and methodological quality of study designs. The proposed assessment therefore does not overestimate the results of deficiently implemented meta-analyses and controlled studies.

Criticism was also levelled at evidence-based practice for neglecting qualitative data and other form of non-scientific information e.g. local traditional knowledge (Adams and Sandbrook, 2013). Some questions can be answered only with qualitative approaches and evidence-based practice does not exclude them (Haddaway and Pullin, 2013; Collins *et al.*, 2014). In the quality checklist not all questions apply equally to quantitative and qualitative research, but this does not bear on the ranking. More reflection and responses to criticism of evidence-based practice can be found in Mullen and Streiner (2004); Sutherland *et al.* (2004, 2005); Haddaway and Pullin (2013).

Despite the criticism raised against evidence-based practice the benefits are clear (Walsh *et al.*, 2014). Statements and recommendations should be based on the best current scientific knowledge, and integrated with expertise, practical experience and the local context. Rating the strength of evidence matters as it allows clarification of the certainty of research results and, thus, of conclusions, decisions, or recommendations drawn from that research (Lohr, 2004).

Wrong decisions are particularly problematic if studies providing strong evidence were available but ignored. Child mortality from sudden infant death syndrome was unnecessarily high for decades due to recommendations ignoring the stronger evidence that was already available at that time (Gilbert *et al.*, 2005). Especially on topics with contradicting opinions - including numerous examples in ecosystem services and conservation, it is important to continuously summarise and update the available evidence.

It is clear that evidence-based practice in environmental management concerns scientists as well as decision-makers and the general public. In the interest of responsible use of environmental resources and processes, we strongly encourage embracing evidence-based practice as a paradigm for all research contributing to environmental management.

## Acknowledgements

We thank Andrew Pullin, Sven Lautenbach and Ian Bateman for valuable comments on earlier versions of the manuscript. This work was supported by the 7th framework programme of the European Commission in the project OPERAs (grant number 308393, www.operas-project.eu).

